# LUMIC: Latent diffUsion for Multiplexed Images of Cells

**DOI:** 10.1101/2024.11.06.622339

**Authors:** Albert Hung, Charles J. Zhang, Jonathan Z. Sexton, Matthew J. O’Meara, Joshua D. Welch

**Affiliations:** Department of Computer Science and Engineering, University of Michigan, Ann Arbor, USA; Computer Science and Artificial Intelligence Laboratory, Massachusetts Institute of Technology, Cambridge, USA; Department of Computational Medicine and Bioinformatics, University of Michigan, Ann Arbor, USA; Department of Medicinal Chemistry, University of Michigan, Ann Arbor, USA; U-M Center for Drug Repurposing, University of Michigan, Ann Arbor, USA

## Abstract

The rapid advancement of high-content, single-cell technologies like robotic confocal microscopy with multiplexed dyes (morphological profiling) can be leveraged to reveal fundamental biology, ranging from microbial and abiotic stress to organ development. Specifically, heterogeneous cell systems can be perturbed genetically or with chemical treatments to allow for inference of causal mechanisms. An exciting strategy to navigate the high-dimensional space of possible perturbation and cell type combinations is to use generative models as priors to anticipate high-content outcomes in order to design informative experiments. Towards this goal, we present the Latent diffUsion for Multiplexed Images of Cells (LUMIC) framework that can generate high quality and high fidelity images of cells. LUMIC combines diffusion models with DINO (self-Distillation with NO labels), a vision-transformer based, self-supervised method that can be trained on images to learn feature embeddings, and HGraph2Graph, a hierarchical graph encoder-decoder to represent chemicals. To demonstrate the ability of LUMIC to generalize across cell lines and treatments, we apply it to a dataset of *∼*27,000 images of two cell lines treated with 306 chemicals and stained with three dyes from the JUMP Pilot dataset and a newly-generated dataset of *∼*3,000 images of five cell lines treated with 61 chemicals and stained with three dyes. To quantify prediction quality, we evaluate the DINO embeddings, Kernel Inception Distance (KID) score, and recovery of morphological feature distributions. LUMIC significantly outperforms previous methods and generates realistic out-of-sample images of cells across unseen compounds and cell types.

## 1 Introduction

High-content imaging assays have revolutionized the ability to observe and analyze the morphological impact of a wide variety of drugs on various different cell types. For example, the Cell Painting assay uses six fluorescent dyes imaged across five different channels to capture cell phenotypes[1], and has already facilitated the identification of drug targets and mechanisms of action[2]. Although morphological profiling assays are cost-effective and straightforward methods that only require commonly available laboratory equipment, the number of possible cell type and chemical compound combinations is infeasible to explore experimentally.

At the same time, substantial progress has been made in generative machine learning, enabling conditional image, video, and text generation[3][4][5]. Deep generative models, such as Generative Adversarial Networks (GANs), normalizing flows, and diffusion models, have recently garnered attention for their ability to generate high-quality and fidelity samples[6][7][8]. Specifically, diffusion models have become popular because of their training stability, ease of guidance, and state-of-the-art performance in image generation tasks. Diffusion models outperform GANs without requiring specific architectural and optimization choices to prevent mode collapse and stable training[8]. During training, a data sample *x*_0_ is slowly corrupted through a *T* step forward Markov chain to create noised samples *x*_1_, *x*_2_, …, *x*_*T*_. Crucially, if the noise follows a Gaussian distribution, then since the sum of Gaussians is also a Gaussian, the noised sample *x*_*t*_ at timestep *t* can be computed efficiently by

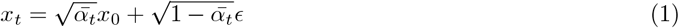

where 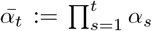 and *α*_*t*_ is the scaling factor controlled by a variance scheduler, and *ϵ* ∼ 𝒩 (0, *I*) is Gaussian noise. Then, to learn the reverse process, a neural network *ϵ*_*θ*_ parameterized by *θ* can be trained to predict the noise at each timestep using an *L*_2_ loss where *t* is the discrete uniform distribution between 1 and T:

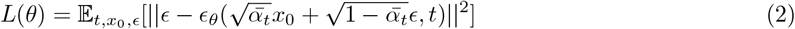

Finally, after the model has been trained, the reverse process can be used to generate new samples by sampling *x*_*T*_ ∼ 𝒩 (0, 1) and iteratively denoising from *T* to 1 using the following equation

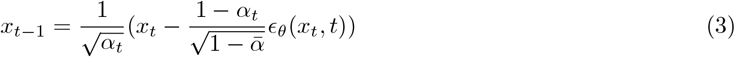

To accelerate inference by generating high quality samples in fewer steps, the Denoising Diffusion Implicit Model (DDIM)[9] can be used instead to sample at a subsequence of the original time schedule, *τ* = [*τ*_1_, *τ*_2_, …, *τ*_*S*_] where *S* < *T*, and is formulated as

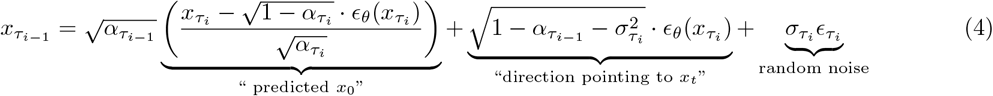

and

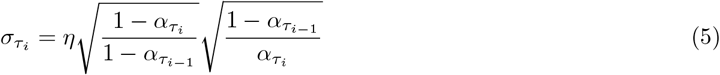

where *η∈* [0, 1] is a hyperparameter that interpolates between 0 to make sampling deterministic and 1 for standard denoising diffusion probabilistic model(DDPM) sampling (Eq. 3).

To enforce conditioning information ***c***, such as class labels or latent representations, classifier-free guidance can be used[10]. This is particularly useful in predicting the outcomes of perturbation experiments, where conditional generation allows for fine-tune control over the desired cell line and chemical interaction by steering the model towards the desired output. The key idea is that given paired data (*x*_0_, *c*), the model learns both a conditional and an unconditional model by randomly dropping out the label during training. Specifically, at time step *t*, the learned noise is defined by

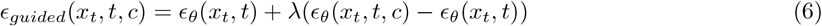

where *ϵ*_*θ*_(*x*_*t*_, *t*) is the noise prediction without conditioning information, *ϵ*_*θ*_(*x*_*t*_, *t, c*) is the noise prediction with conditioning, and λ is the guidance weight.

With the rise of generative models and the curation of large biological datasets, it is now feasible and promising to train generative models on biological data. For example, the JUMP dataset contains 116k chemical perturbations[2]–analogous to the massive datasets used to train generative models in the natural language and vision space. However, no previous methods has attempted generation of multiple cell types, limiting downstream applicability. Mol2Image presents a flow-based model to generate Cell Painting images conditional on a graph neural network chemical embedding [11]. Various other authors have also attempted to use GANs to generate Cell Painting images, including IMage Perturbation Autoencoder(IMPA)[12], GANs as Robust Adversarial Perturbation Encoders(GRAPE)[13], Deep Convolutional GAN (DCGAN)[14], CP2Image[15], and Interventional Style Transfer(IST)[16]. IMPA adopts a conditional GAN to style transfer perturbations onto U2OS cells, treating cells as content and compounds as the style. GRAPE builds on IMPA to leverage style transfer using generative modeling to learn representations of genetic perturbations. The authors of [14] use DCGAN, a GAN in which the pooling layers are replaced with strided convolutions, to generate high-content microscopy images, but the scope of their study was limited, as it was only trained and evaluated only on ten compounds. CP2Image trains a GAN conditional on CellProfiler representations to generate images but using CellProfiler features lacks the flexibility of other methods to directly control the chemical compound used to treat a cell. IST utilizes an encoder-decoder U-Net architecture as part of their network to generate training data with mitigated spurious correlations. Diffusion models, specifically image-to-image diffusion models, have also been used to generate fluorescent microscopy images, including PhenDiff[17], and guided-I2I[18]. PhenDiff attempts to “translate” between perturbations conditional on class, and guided-I2I converts brightfield images to fluorescent microscopy images conditional on a chemical class label; however, these methods are unable to predict the responses of chemicals not seen during training.

LUMIC adopts a standard DDPM pipeline and is not only able to beat existing methods on single cell line generation tasks but also removes the limitation of single cell line generation, allowing for the controllable prediction of multiple cell line and chemical compound interactions. Specifically, our key innovation is to model how perturbing a specific type of cell (specified as an image of the control cell well) changes its morphology. Analogous to text-to-image approaches such as DALL-E 2 and stable diffusion, we use diffusion in the image embedding space (latent diffusion) to learn the context-specific effect of a compound[19][20]. This allows us to predict how either seen or unseen compounds will affect either seen or unseen cell lines. To evaluate our approach of “transferring” perturbations onto a new cell line, we performed laboratory experiments to generate a new Cell Painting dataset featuring the same treatments across multiple cell lines. Biological and computational evaluations demonstrate that LUMIC can meaningfully predict the effects of chemical treatment for both unseen compounds and unseen cell types.

## 2 Methods

### 2.1 LUMIC: Latent Diffusion for Multiplexed Images of Cells

The LUMIC pipeline is built on the following intuition: a particular cell type has a baseline unperturbed morphology with some variation between cells. Treating these cells with a particular small molecule shifts the internal states of the cells and ultimately results in a morphological change (again with some possible variation among cells). We further expect that similar cells exposed to similar molecules will undergo similar changes (for appropriate definitions of “similar”).

Following this intuition, LUMIC uses conditional diffusion in the latent image space to predict the embedding of the perturbed cell from the embeddings of the unperturbed cell and the small molecule. To do this, we need a way of embedding both control and perturbed images; a way to embed small molecules; a network to predict perturbed image embedding; and a way to predict a high-resolution perturbed image from its embedding in latent space.

We use a pre-trained DINO model[21] to embed images and a pre-trained Hgraph2Graph model[22] to embed small molecules. We train three diffusion models (**Fig. 1**). The first diffusion model generates the image embedding of a particular control cell image treated with a particular molecule. The second diffusion model generates a low-resolution 64 × 64 image from an image embedding. The last diffusion model generates a high-resolution 256 × 256 image from a 64 × 64 image. A key advantage of separating embedding generation and image generation is that the image embedding can be useful for downstream tasks where the actual image is not required, such as understanding biological variation[23]. Separating the image generation (low-resolution and high-resolution) process into to separate models instead of a single high-resolution model helps to ensure high-quality samples and efficient training and sampling[24].

**Figure 1:**
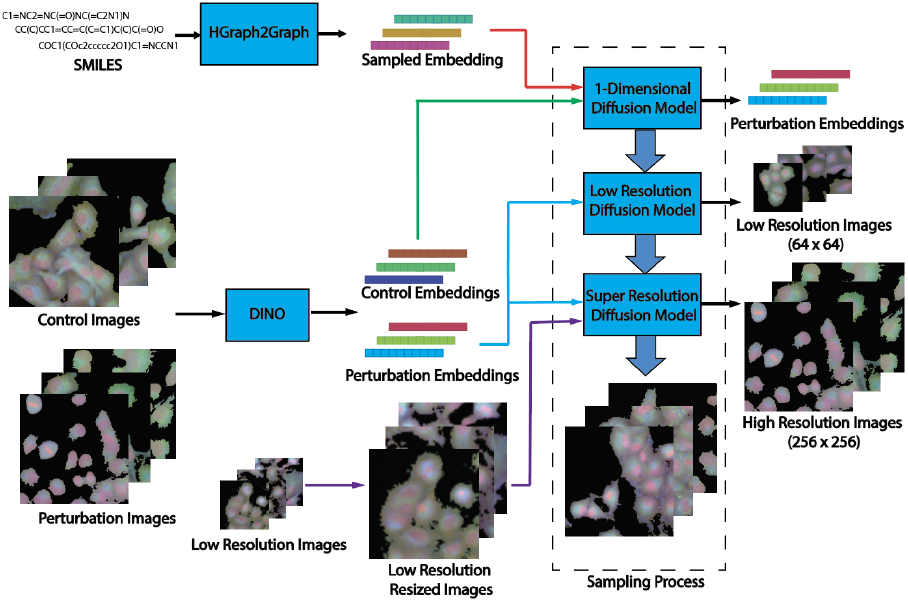
Architecture Diagram of Diffusion Pipeline

**Figure 2:**
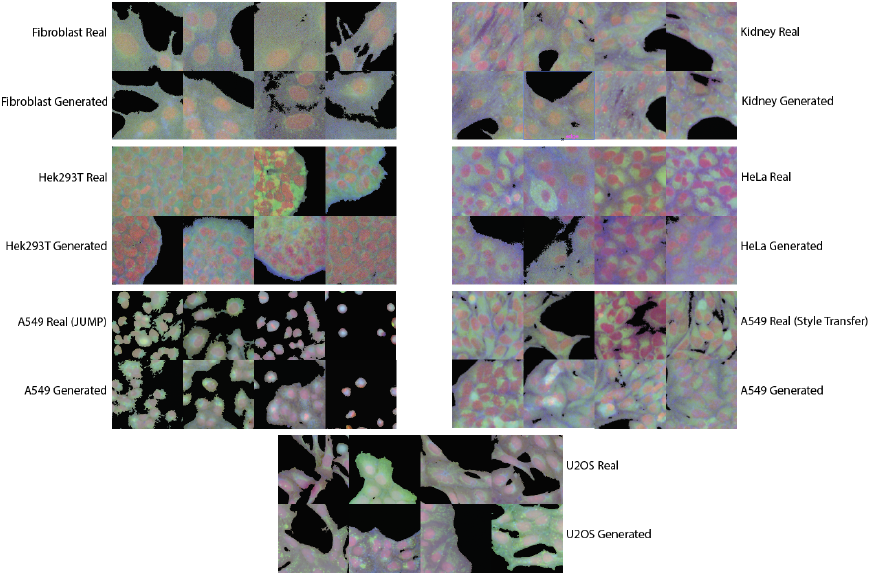
Real and generated samples separated by cell type

#### 2.1.1 Embedding Control and Perturbed Images with DINO

To learn image feature embeddings that capture cell line information, we trained a DINO (self-Distillation with NO labels) model, which is a vision-transformer that uses a self-supervised loss[21]. DINO has shown to be effective in learning representations of cellular morphology directly from images[23] and outperforms other self-supervised learning methods, including SymCLR and MAE, as well as computer vision based feature embeddings from CellProfiler in downstream tasks[25]. We trained our DINO model on *∼* 27,000 images from the JUMP Pilot subset and *∼* 3,000 images for our style transfer dataset. During training, we used additional preprocessing steps and transformations including taking random 338 crops and resizing them to 224 and randomly dropping out channels[23]. During sampling, we padded images by 50 pixels on each side to help DINO focus on the content rather than gaps between cells, since some of the cells cultured for the style transfer dataset were partially to fully confluent, as shown by visualizing the attention maps.

#### 2.1.2 Embedding Small Molecules with HGraph2Graph

To encode chemical information, we trained an HGraph2Graph model, which is a hierarchical graph encoder-decoder that utilizes structural motifs as building blocks to encode SMILES[22], on all 306 chemical compounds from the JUMP Pilot dataset, 61 compounds from our style transfer dataset, and 250k compounds randomly sampled from the ZINC dataset by[26]. During training of LUMIC, the SMILES are passed into Hgraph2Graph, and a 128-dimensional vector is sampled from the latent space. The remaining 256 dimensions are padded with zeros.

#### 2.1.3 Image Embedding Diffusion Model

To predict image embeddings with diffusion, we modified the standard U-Net architecture to use 1-dimensional instead of 2-dimensional convolutions over the image embedding (a vector) instead of an image (a matrix). Given a control image of a cell line, we first take a 256 × 256 random crop of the image and encode it into a 384-dimensional feature embedding using DINO. Then a compound (SMILES) is encoded using HGraph2Graph and a chemical latent is sampled. The model then outputs the visual embedding of the targeted interaction (that cell type treated with that compound). This diffusion model employs linear attention and self-attention to learn conditional information. We trained this model using the Adam optimizer with a learning rate of 5e-4 and a batch size of 64 for 24 hours, totaling 75,000 steps.

#### 2.1.4 Low-Resolution Diffusion Model

The low resolution diffusion model takes the visual embeddings (from DINO) and decodes them into their respective image. The model takes random (256 × 256) crops of images, encodes them, and learns to generate the (64 × 64) low resolution image based on the embedding. This model follows the efficient U-Net architecture proposed in[3] and uses cross attention at each layer to capture the conditioning information. We trained this model using the Adam optimizer with a learning rate of 5e-5 and a batch size of 64 for 24 hours totaling 150,000 steps.

#### 2.1.5 Super-Resolution Diffusion Model

The super resolution diffusion model takes the visual embeddings and the low resolution (64 × 64) image and generates the corresponding high resolution (256 × 256) image. The model is trained by taking 256 × 256 random crops and inputting the low resolution (resizing the 256 × 256 crop to 64 × 64 and then resizing it back to 256 × 256) version as well as the encoded version (passing the 256 × 256 crop into DINO). The model architecture is consistent with the efficient U-Net architecture used in[3] with linear attention. We trained this model using the Adam optimizer with a learning rate of 5e-5 and a batch size of 33 for ten days, totaling 1,000,000 steps across 3 A40 GPUs.

#### 2.1.6 Training Details

All three diffusion models were trained independently. During training of the diffusion models, random crops were resampled up to 50 times if the percentage of black pixels exceeded 30% of the entire image. We used random crops instead of single-cell crops in order to capture how the cells grow together and to capture how a chemical may impact this growth. Moreover, if the growth pattern of a cell and compound interaction is very sparse, we want our generated images to reflect this. Then, the images were randomly flipped horizontally (p = 0.5), channels were randomly dropped out (p = 0.2), and finally images were padded by 50 before being passed into DINO. All models were trained on single NVidia A40 single precision GPUs (unless otherwise mentioned) until the visual quality of the images reached a sufficient level. All diffusion models were also trained using exponential moving average (EMA), averaging over the past ten training weights, which has shown to improve sampling quality[27]. The EMA model was also used during sampling. A cosine noise schedule with 1,000 timesteps was used during training of all diffusion models, while a linear noise schedule with 1,000 timesteps was used for the low resolution noise schedule for the super-resolution model.

### 2.2 Datasets

The JUMP Pilot Target 1 dataset is a subset of the JUMP Cell Painting dataset and consists of 306 different chemical perturbations on U2OS cells (derived from a bone cancer of female donor) and A549 cells (derived from a lung cancer of a male donor) at 2 different time points (24 and 48 hours). We used only the 24 hour timepoint, resulting in 4 plates per cell line and 27,702 images total. Additionally, pairs of compounds within this dataset are known to target the same protein encoded by a given gene[28]. We split the data based on the gene that they target: a total of 30 compounds are held out, 16 compounds target genes not seen during training (8 genes), which we call unseen genes, and 14 compounds target genes seen during training, which we call seen genes. Three of the five fluorescent channels–nucleus (Hoechst; DNA), actin cytoskeleton/golgi/plasma membrane (phalloidin; AGP), and mitochondria (MitoTracker; Mito)–were stacked to form an RGB image for compatibility with the standard DINO architecture.

We also generated a new experimental dataset, which we refer to as the “style transfer dataset”. To generate the data, multiple cell lines were treated with the same panel of compounds. Each compound was dissolved in DMSO at a concentration of 2 mM and distributed into an Echo Qualified 384-Well Polypropylene Microplates (Beckman Coulter, 001-14615). Using an Echo 655 Liquid Handler, 250 nL of each compound was dispensed into 384-well PhenoPlates (Revvity, 6057302) for a final concentration of 10 µM at a final volume of 50 µL/well.

3T3, A549, HEL293T, HeLa, and RPTE cells were grown to 90% confluency in DMEM/F12 (ThermoFisher Scientific, 11320033) supplemented with 10% FBS (ThermoFisher Scientific, A5256701) in T75 flasks. Media was then aspirated and cells were washed with 10 mL of PBS (ThermoFisher Scientific, 10010023) before lifting with 1 mL of 0.25% Trypsin-EDTA (ThermoFisher Scientific, 25200056). After 5 minutes of incubation, 10 mL of the 10% FBS DMEM/F12 was added into each flask to remove cells. Cells were spun down at 300 x g for 5 minutes to pellet and resuspended in fresh 10% FBS DMEM/F12 before counting by hemocytometer. Cells were diluted to a final concentration of 120,000 cells/mL and 50 µL was dispensed into each well for a final seeding density of 6,000 cells/well.

Plates were incubated at 37°*C* for 24 hours before fixing and staining using the same stains as the JUMP Pilot data (DNA, AGP, Mito) in a CellPainting style assay[1]. We used a Yokogawa CQ1 High-Content Imaging system to image multiple fields of view for each well, for a total of 3,168 images. We randomly select 10 compounds to hold out across all 5 cell lines and hold out HeLa completely (except control images) during training to make up the test set.

For both datasets, the images were normalized using sklearn’s Quantile Transformation with respect to the controls[29]. For the style transfer dataset, Cellpose was used to identify the nuclei of the cells and create a binary mask[30], and CellProfiler was used to segment the images to remove any residual dye in the background[31].

### 2.3 Evaluation

Approximately 1,000 images were generated for each class using the entire generative modeling pipeline (generating the embedding, decoding that embedding into the 64 × 64 image, and generating the 256 × 256 image from the generated embedding and low resolution generated image shown in the vertical dashed box in **Fig. 1**). For all 3 models, DDIM sampling (*η* = 0) was used with 250 steps, which was where the decrease in KID between differing sampling steps became less drastic.

## 3 Results

### 3.1 LUMIC Generates Realistic Image Embeddings that Preserve Cell Type and Treatment Semantics

To qualitatively evaluate the generated representations, we plotted the UMAP embeddings of the real and generated DINO features colored by cell type and compounds (**Fig. 3**). Both the real and generated DINO embeddings cluster well by cell type, and the generated embeddings align with their respective ground truth clusters, suggesting that the generated embeddings contain corresponding cell type specific information. An additional cluster isolated from the rest contains a mixture of cell types. The images in this cluster largely consist of cell-free black backgrounds because of how we randomly cropped images.

**Figure 3:**
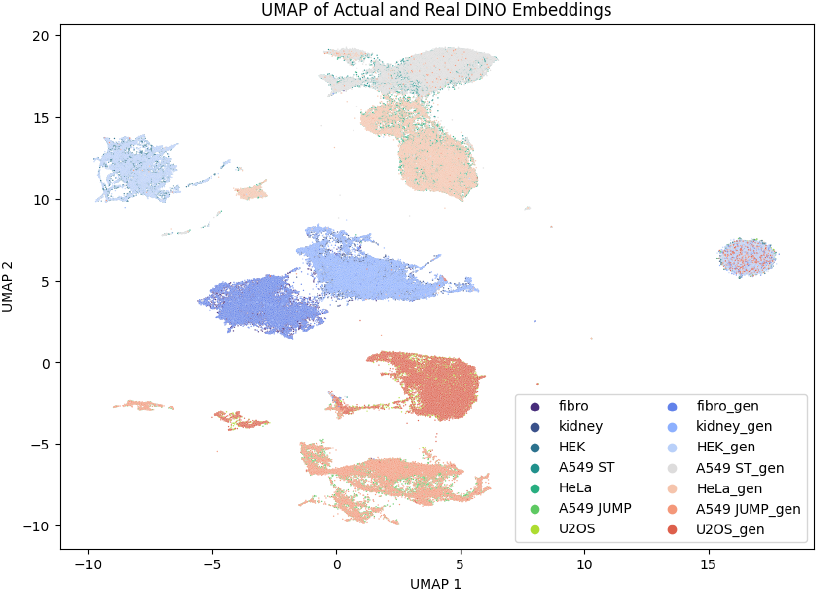
UMAPs of real and generated embeddings grouped by cell type

The real and generated image embeddings also reflect differences among chemical treatments to some extent **Fig. 4**). The fibroblast cells (**4A** and **4B**) separate somewhat less clearly than the HeLa cells (**Fig. 4C** and **4D**), possibly reflecting cell type differences in the magnitude of morphology change. Nevertheless, treatment differences among cells are apparent in both cases. In particular, images of HeLa cells treated with a given compound tend to cluster with each other and apart from images of cells treated with a different compound. This phenomenon is also seen in the JUMP datasets. A549 cells (**Sup. Fig. A1**) show relatively less separation among treatments compared with U2OS cells (**Sup. Fig. A2**), which have almost layer-like subclusters. Nevertheless, the UMAPs of the generated embedding generally reflect the shape and distribution of their intended cell type and compound. This indicates that both the real and generated DINO image embeddings likely do reflect differences in cell type and chemical treatment for both seen and unseen cell types and chemical compounds.

**Figure 4:**
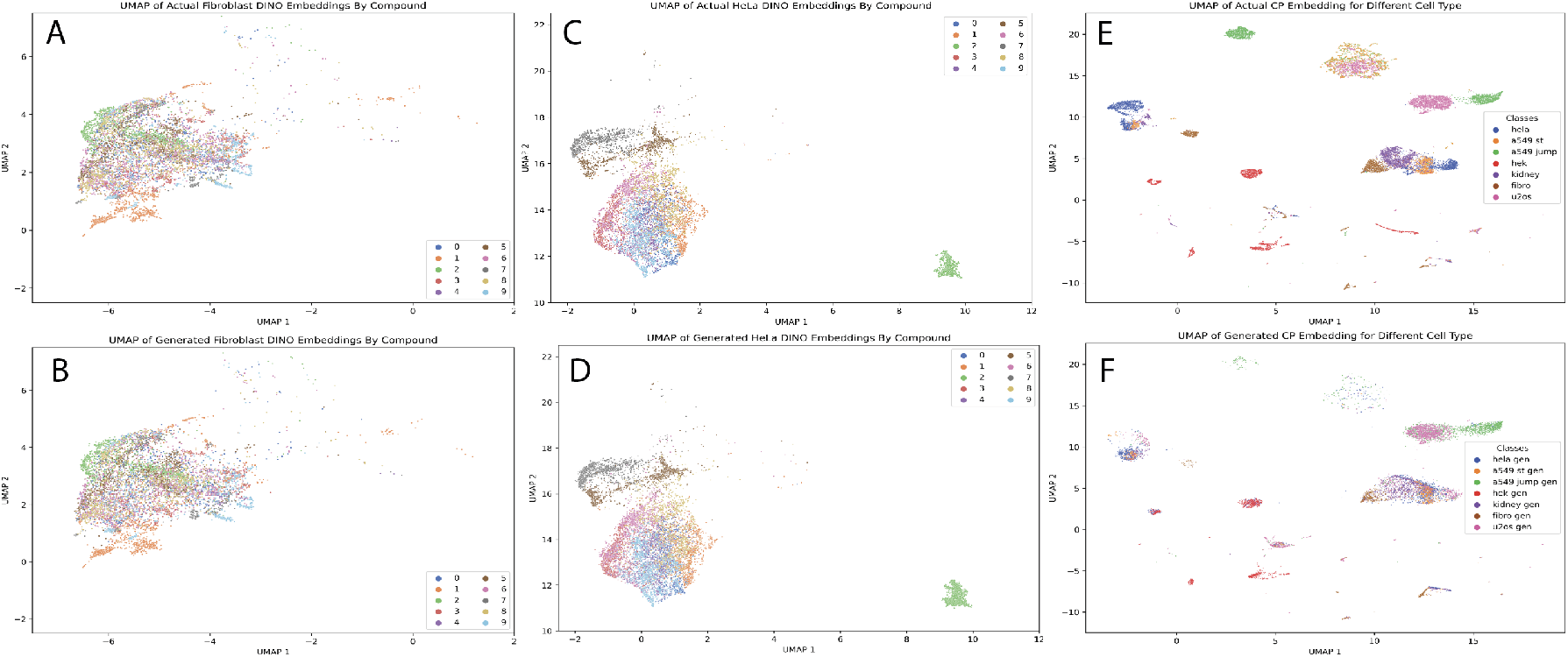
UMAPs of real and generated embeddings grouped by compound

To quantitatively evaluate whether the generated embeddings reflect cell type semantics, we trained MultiLayer Perceptron (MLP) and k-Nearest-Neighbor (KNN) models to predict cell type from DINO embedding. We reasoned that if a classifier could accurately identify the cell type of a generated embedding, then LUMIC is correctly obeying the cell type conditioning. All model training was done using scikit-learn and a validation set of 20% (of actual embeddings) was held out during training[29]. Reassuringly, the performance of the MLP and the KNN models is very similar on the held out set of real image embeddings and the set of generated embeddings from control cells of each type (**Table 1**).

**Table 1:** Accuracy of Cell Type and Treatment Classification using DINO Embeddings.

|  | Control Images |  | Unseen Compounds |  |
| --- | --- | --- | --- | --- |
|  | Real | Generated | Real | Generated |
| MLP | 0.925 | 0.916 | 0.929 | 0.918 |
| KNN | 0.873 | 0.877 | 0.912 | 0.915 |

We next used a similar strategy to evaluate whether embeddings generated by LUMIC preserve treatment semantics. The MLP and KNN classifiers trained to identify held out treatment from embeddings of real cells were still able to identify the held out treatment of generated image embeddings (**Table 3**). The small drop in classification performance between real and generated embeddings indicates that the embeddings generated by LUMIC do indeed reflect the differences among images from different treatments, even for treatments not seen during LUMIC training.

**Table 2:**
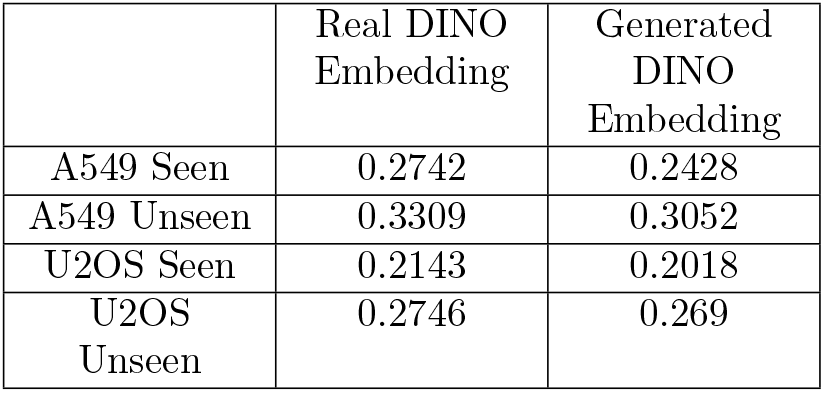
Accuracy of Gene Classification using KNN.

|  | Real DINO Embedding | Generated DINO Embedding |
| --- | --- | --- |
| A549 Seen | 0.2742 | 0.2428 |
| A549 Unseen | 0.3309 | 0.3052 |
| U2OS Seen | 0.2143 | 0.2018 |
| U2OS Unseen | 0.2746 | 0.269 |

**Table 3:** Accuracy of Compound Classification (test set) using DINO Embeddings.

|  | MLP |  | KNN |  |
| --- | --- | --- | --- | --- |
|  | Real | Generated | Real | Generated |
| A549 | 0.859 | 0.721 | 0.741 | 0.729 |
| Kidney | 0.81 | 0.696 | 0.651 | 0.647 |
| Fibroblast | 0.724 | 0.609 | 0.571 | 0.574 |
| HEK293T | 0.554 | 0.47 | 0.406 | 0.409 |
| HeLa | 0.869 | 0.724 | 0.729 | 0.735 |

To further evaluate the quality of the generated embeddings, we trained a KNN classifier to identify the target gene of the compound used in each treatment (recall that the JUMP Pilot dataset profiled a set of compounds with a total of 146 known target genes). We trained the classifier on the DINO embeddings of real images, then ran this classifier on image embeddings generated by LUMIC to quantify how well the embeddings retained signatures of the gene targeted by the compound. A high classifier score would indicate that the generated embeddings do indeed preserve the differences between images of cells treated with different compounds; the classifier performance on real embeddings represents an upper bound on generation performance. As seen in **Table 2**, the classifier performance on real and generated embeddings is similar, indicating that the model does indeed generate embeddings that distinguish among drug treatments. The classifier accuracy on generated embeddings relative to real embeddings is lower for A549 cells than U2OS cells. This likely reflects the observation discussed above–DINO representations of real A549 cells do not cluster as well by drug treatment as real U2OS cells. Nevertheless, these results indicate that we are able to meaningfully encode the targeted gene in the generated embeddings, capturing the underlying biology of the interactions.

### 3.2 LUMIC Generates More Realistic Morphology Images than Previous Approaches

We next compared LUMIC against previous approaches for morphology image generation. Because previous approaches cannot generate images for unseen cell types and/or molecules, we chose to evaluate unconditional single cell line generation. To do this, we calculated the Kernel Inception Distance (KID) between the real and generated images. KID is the squared maximum mean discrepancy between the distributions of image representations from the Inception V3 network. Smaller KIDs are better, indicating smaller distance between true and generated image distributions. LUMIC outperforms the other methods, achieving a significantly lower KID (**Table 4**). Thus, our model is able to capture accurate morphology and growth patterns with fidelity that surpasses the existing state of the art for single cell type generation.

**Table 4:** KID of Unconditional U2OS generation using Random Crops during Training.

|  | KID |
| --- | --- |
| LUMIC | **0.0154** |
| PhenDiff | 0.3313 |
| IMPA | 0.1727 |

**Table 5:** KID of real vs. generated cell lines. Rows represent real images from each cell line; columns are generated images. Lower KID is better. Low on-diagonal KID and high off-diagonal KID indicate that LUMIC beats the baseline.

|  | A549 | HEK293T | Fibroblast | Kidney | HeLa |
| --- | --- | --- | --- | --- | --- |
| A549 | **0.0132** | 0.214 | 0.082 | 0.0781 | 0.0425 |
| HEK293T | 0.2558 | **0.0069** | 0.134 | 0.1089 | 0.02225 |
| Fibroblast | 0.0819 | 0.0936 | **0.0114** | 0.0672 | 0.0641 |
| Kidney | 0.0533 | 0.1608 | 0.035 | **0.0158** | 0.0641 |
| HeLa | 0.0488 | 0.2042 | 0.1006 | 0.1089 | **0.0346** |

### 3.3 Generated Images from Unseen Cell Types and Treatments Outperform Baseline Model

We next assessed whether LUMIC can meaningfully predict images from unseen cell types and unseen treatments by comparing it with a baseline. We reasoned that, in order to be useful, the model predictions for a given cell type must be more similar to real images of the same cell type than real images of a different cell type. Similarly, for a cell type/treatment combination, the generated images must be more similar to the real cell type/treatment combination than to untreated control cell images. This baseline is not trivial, because it requires the generative model to preserve both the overall characteristics of morphology images and the semantics of a particular cell line and/or treatment. Reassuringly, LUMIC beats this baseline for image generation conditional on cell type by a large margin (**Fig. 5**), indicating that LUMIC both generates realistic images and obeys the semantics of cell type conditions.

**Figure 5:**
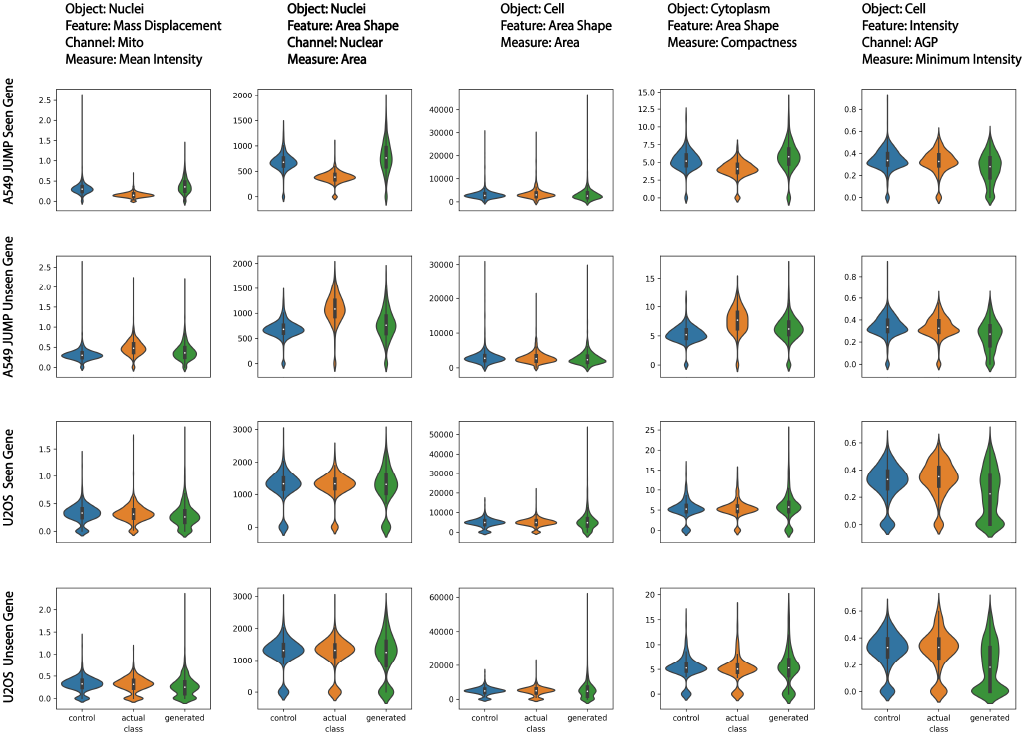
Violin Plot of the most distinguishing features for each class from SHAP

We next assessed LUMIC images generated from specified cell type/treatment combinations. We investigated three prediction tasks of increasing difficulty: (1) the cell type is in the training data but all images of a treatment are held out (seen cell type, unseen treatment; **Table 7**); (2) all treatments of a given cell type are held out, but the treatments are observed for other cell types (unseen cell type, seen treatment; **Table 8**); and (3) a cell type/treatment combination is predicted when neither the cell type nor the treatment have been observed during training (unseen cell type, unseen treatment; **Table 8**). For the JUMP dataset, we also investigated the difference between treatments with compounds targeting seen or unseen genes (**Table 7**). Remarkably, LUMIC outperforms the baseline across all three prediction tasks. Specifically, for the JUMP dataset, where compounds are held out that impact seen and unseen genes, LUMIC not only beats the baseline comparison against control images but even performs better on compounds that impact unseen genes, suggesting the ability to remain accurate on out of distribution chemical perturbations (**Table 6**). When looking at the seen cell lines in the style transfer dataset, LUMIC is able to predict effects of unseen compounds on multiple seen cell types, suggesting its applicability in downstream use cases to expand the scope of existing screens (**Table 7**). Finally, when we held out all HeLa cell treatments, LUMIC was able to predict both seen and unseen compounds on an unseen cell line better than baseline, suggesting that it is able to create entirely new screens provided only control images (**Table 8**). Notably, the generation quality for predictions of the unseen cell type was worse, with the within class KID being much closer to baseline KID values than in the easier generation tasks.

**Table 6:** KID of seen and unseen targeted genes on seen cell lines.

| Cell Line | Corresponding Class | Control Images of Same Cell Type |
| --- | --- | --- |
| U2OS Seen Gene | **0.0289** | 0.0404 |
| U2OS Unseen Gene | **0.0287** | 0.0539 |
| A549 Seen Gene | **0.0162** | 0.0276 |
| A549 Unseen Gene | **0.0138** | 0.0345 |

**Table 7:** KID of unseen compounds on various seen (A549, Fibroblast, Kidney, HEK293T) cell lines. We report the average KID across cell types for compounds.

| Cell Line | Corresponding Class | Control Images of Same Cell Type |
| --- | --- | --- |
| A549 | **0.0512** | 0.1133 |
| Fibroblast | **0.0254** | 0.1072 |
| Kidney | **0.0173** | 0.0349 |
| HEK293T | **0.0094** | 0.0446 |

**Table 8:** KID of seen and unseen compounds on HeLa.

| Compounds | Corresponding Class | Control Images of Same Cell Type |
| --- | --- | --- |
| Seen Compounds | **0.0748** | 0.0788 |
| Unseen Compounds | **0.0742** | 0.0748 |

### 3.4 Intensity Features Show Largest Difference Between Real and Generated Images

Cell morphology images from the Cell Painting dyes are often analyzed using hand-crafted features, such as those from the CellProfiler pipeline. These features capture various aspects of cell and nuclear shape and size, as well as dye intensity. To further investigate how well our generated images reflect biological properties of cells, we ran CellProfiler to segment cells and extract features. We then removed highly correlated features (*>* 0.9) and features with low variance (*<* 0.01). Then, we trained MLP and KNN models to classify cell type from CellProfiler features of real cells, and evaluated the classifier on CellProfiler features of generated cell images (**Table 9**). The classifier was much less accurate at identifying the cell types of generated images from their CellProfiler features, indicating some sort of distribution shift.

**Table 9:** Accuracy of Cell Type Classification using CellProfiler Features.

|  | Train | Validation | Generated |
| --- | --- | --- | --- |
| MLP | 0.72 | 0.735 | 0.472 |
| KNN | 0.527 | 0.507 | 0.349 |
| LR | 0.844 | 0.849 | 0.509 |

To understand which features show the largest differences between real and generated images, we trained a random forest (per cell type) with 100 decision trees to classify real and generated images (**Table 10**). We then used SHapley Additive exPlanations (SHAP) to identify the most important features per class[32]. The top 5 most distinguishing features from the 10 types of CellProfiler features are shown in **Table 11**. All of these features are a result of intensity based metrics, with the area shape measurements being calculated after manual intensity based thresholding to identify the nucleus. A UMAP of the real and generated CellProfiler indicates that they do not match up as well as the real and generated DINO embeddings as seen in **Fig. 4E** and **4F**. Specifically, the bottom most green cluster that is A549 JUMP compounds that is present in the actual embeddings but not in the generated ones and the noticeable blue cluster of HeLa on the right side, which is again present in the actual embedding UMAP but not on the generated ones. This suggests that the biological features extracted from the generated images do not accurately match those extracted from the actual images. Moreover, when looking at the violin plots from the most differentiating factors in **Fig. 5**, there is a lot of variation between cell lines and between features. For example, the generated distribution for the `Cytoplasm Areashape Compactness` feature for HeLa drastically differs in shape compared to both the controls and actual images, whereas for HEK, the generated distribution for `Nuclei Intensity MassDisplacement Mito` is more similar to the actual class distribution than the controls. This variance suggests that there are still areas for this model to improve in terms of biological realism. However, since all of these features that allow the RF to distinguish between real vs generated are intensity based or are easily impacted by varying intensity, we hypothesize that the large variance in generated features reflects a drastic variance in intensity that our model generates. This may make it difficult to accurately calculate CellProfiler features using manual thresholding. Thus, this analysis suggests that there is room for future improvement by better calibrating the distribution of generated intensity values.

**Table 10:**
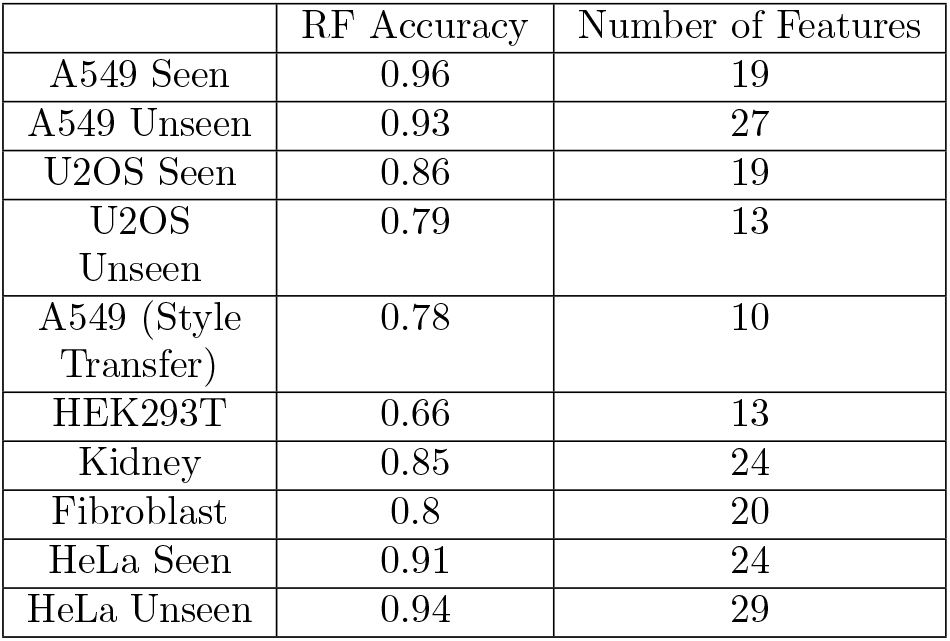
RF Accuracy and Number of Features to Classify Between Real and Generated CellProfiler Features.

|  | RF Accuracy | Number of Features |
| --- | --- | --- |
| A549 Seen | 0.96 | 19 |
| A549 Unseen | 0.93 | 27 |
| U2OS Seen | 0.86 | 19 |
| U2OS Unseen | 0.79 | 13 |
| A549 (Style Transfer) | 0.78 | 10 |
| HEK293T | 0.66 | 13 |
| Kidney | 0.85 | 24 |
| Fibroblast | 0.8 | 20 |
| HeLa Seen | 0.91 | 24 |
| HeLa Unseen | 0.94 | 29 |

**Table 11:** Accuracy of Cell Type Classification using CellProfiler Features.

| Feature | Appearances within Top 5 Most Influential Features |
| --- | --- |
| Nuclei Intensity<br>MassDisplacement Mito | 7 |
| Nuclei AreaShape Area | 6 |
| Cell AreaShape Area | 5 |
| Cytoplasm AreaShape<br>Compactness | 5 |
| Cell Intensity<br>MinIntensity AGP | 4 |

## 4 Conclusion

In this study, we demonstrate the effectiveness of LUMIC in being able to accurate predict cellular morphologies of seen cell types and unseen compounds, unseen cell types and seen compounds, and unseen cell type and unseen compounds. Heavily influenced by ”style transfer”, we leveraged this idea in our diffusion pipeline to treat chemical perturbations as a ”style” that can be transferred across various different cell lines by first generating a vision embedding of the interaction based on the control image and a chemical embedding, before decoding the vision embedding into a high resolution image. We validate our findings both computationally and biologically, further implicating the possibilities of generative models in facilitating drug discovery and improved cellular understanding.

## 5 Code Availability

The code for our model is available [https://github.com/welch-lab/LUMIC]. The data is available at [https://huggingface.co/datasets/azhung/StyleTransferData]. The model checkpoints are available at [https://huggingface.co/azhung/StyleTransferData/tree/main].

## 6 Acknowledgements

This work was supported by NIH grants R35 GM151129-01 to MJO and R01HG010883 to JDW.

## References

[1] Bray, M.-A. et al. Cell Painting, a high-content image-based assay for morphological profiling using multiplexed fluorescent dyes. Nature protocols 11, 1757–1774 (2016).

[2] Chandrasekaran, S. N. et al. Jump cell painting dataset: morphological impact of 136,000 chemical and genetic perturbations. BioRxiv 2023–03 (2023).

[3] Saharia, C. et al. Photorealistic text-to-image diffusion models with deep language understanding. Advances in Neural Information Processing Systems 35, 36479–36494 (2022).

[4] Ho, J. et al. Imagen video: High definition video generation with diffusion models. arXiv preprint arXiv:2210.02303 (2022).

[5] Achiam, J. et al. GPT-4 technical report. arXiv preprint arXiv:2303.08774 (2023).

[6] Goodfellow, I. et al. Generative adversarial networks. Communications of the ACM 63, 139–144 (2020).

[7] Rezende, D. & Mohamed, S. Variational inference with normalizing flows. In International conference on machine learning, 1530–1538 (PMLR, 2015).

[8] Ho, J., Jain, A. & Abbeel, P. Denoising diffusion probabilistic models. Advances in neural information processing systems 33, 6840–6851 (2020).

[9] Song, J., Meng, C. & Ermon, S. Denoising diffusion implicit models. arXiv preprint arXiv:2010.02502 (2020).

[10] Ho, J. & Salimans, T. Classifier-free diffusion guidance. arXiv preprint arXiv:2207.12598 (2022).

[11] Yang, K. et al. Mol2Image: improved conditional flow models for molecule to image synthesis. In Proceedings of the IEEE/CVF Conference on Computer Vision and Pattern Recognition, 6688–6698 (2021).

[12] Palma, A., Theis, F. J. & Lotfollahi, M. Predicting cell morphological responses to perturbations using generative modeling. bioRxiv 2023–07 (2023).

[13] Bigverdi, M. et al. Gene-level representation learning via interventional style transfer in optical pooled screening. In Proceedings of the IEEE/CVF Conference on Computer Vision and Pattern Recognition, 7921–7931 (2024).

[14] Hussain, S. et al. High-content image generation for drug discovery using generative adversarial networks. Neural Networks 132, 353–363 (2020).

[15] Ji, Y., Cutiongco, M., Yuan, K. et al. CP2Image: Generating high-quality single-cell images using CellProfiler representations. In Medical Imaging with Deep Learning, 274–285 (PMLR, 2024).

[16] Pernice, W. M. et al. Out of distribution generalization via interventional style transfer in single-cell microscopy. In Proceedings of the IEEE/CVF Conference on Computer Vision and Pattern Recognition, 4326–4335 (2023).

[17] Bourou, A. et al. PhenDiff: Revealing invisible phenotypes with conditional diffusion models. arXiv preprint arXiv:2312.08290 (2023).

[18] Cross-Zamirski, J. O. et al. Class-guided image-to-image diffusion: Cell painting from brightfield images with class labels. arXiv preprint arXiv:2303.08863 (2023).

[19] Rombach, R., Blattmann, A., Lorenz, D., Esser, P. & Ommer, B. High-resolution image synthesis with latent diffusion models. In Proceedings of the IEEE/CVF conference on computer vision and pattern recognition, 10684–10695 (2022).

[20] Ramesh, A., Dhariwal, P., Nichol, A., Chu, C. & Chen, M. Hierarchical text-conditional image generation with CLIP latents (2022). 2204.06125.

[21] Caron, M. et al. Emerging properties in self-supervised vision transformers. In Proceedings of the IEEE/CVF international conference on computer vision, 9650–9660 (2021).

[22] Jin, W., Barzilay, R. & Jaakkola, T. Hierarchical generation of molecular graphs using structural motifs. In International conference on machine learning, 4839–4848 (PMLR, 2020).

[23] Doron, M. et al. Unbiased single-cell morphology with self-supervised vision transformers. bioRxiv 2023–06 (2023).

[24] Ho, J. et al. Cascaded diffusion models for high fidelity image generation. Journal of Machine Learning Research 23, 1–33 (2022).

[25] Kim, V. et al. Self-supervision advances morphological profiling by unlocking powerful image representations. BioRxiv 2023–04 (2023).

[26] Kusner, M. J., Paige, B. & Hernandez-Lobato, J. M. Grammar variational autoencoder. In International conference on machine learning, 1945–1954 (PMLR, 2017).

[27] Nichol, A. Q. & Dhariwal, P. Improved denoising diffusion probabilistic models. In International conference on machine learning, 8162–8171 (PMLR, 2021).

[28] Chandrasekaran, S. N. et al. Three million images and morphological profiles of cells treated with matched chemical and genetic perturbations. Nature Methods 1–8 (2024).

[29] Pedregosa, F. et al. Scikit-learn: Machine learning in Python. Journal of Machine Learning Research 12, 2825–2830 (2011).

[30] Stringer, C., Wang, T., Michaelos, M. & Pachitariu, M. Cellpose: a generalist algorithm for cellular segmentation. Nature methods 18, 100–106 (2021).

[31] Stirling, D. R. et al. CellProfiler 4: improvements in speed, utility and usability. BMC bioinformatics 22, 1–11 (2021).

[32] Lundberg, S. A unified approach to interpreting model predictions. arXiv preprint arXiv:1705.07874 (2017).

