## Supplemental Figures for "LUMIC: Latent diffUsion for Multiplexed Images of Cells"

### Supplements

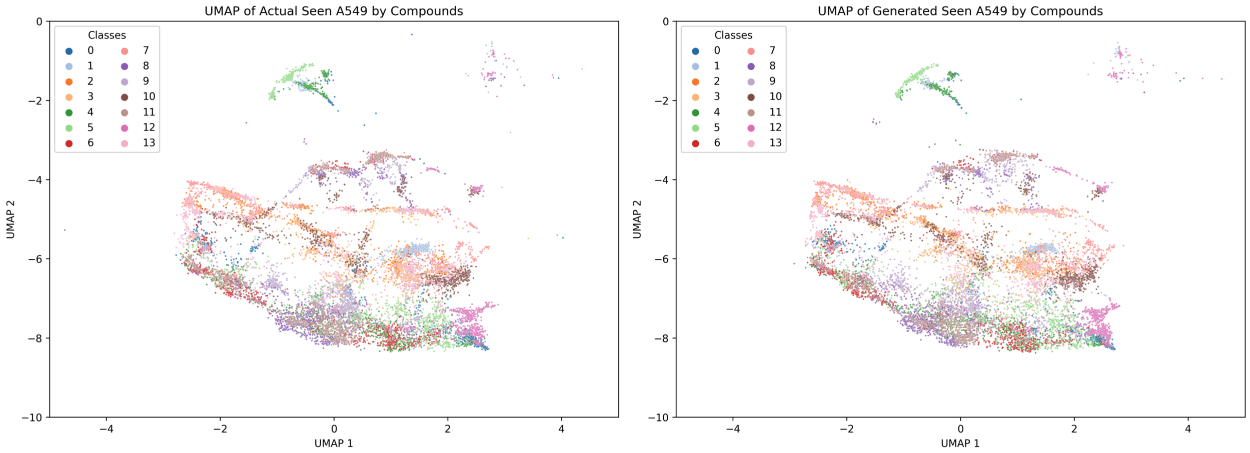


Figure A1: UMAP of real and generated embeddings for seen genes on A549 (JUMP)

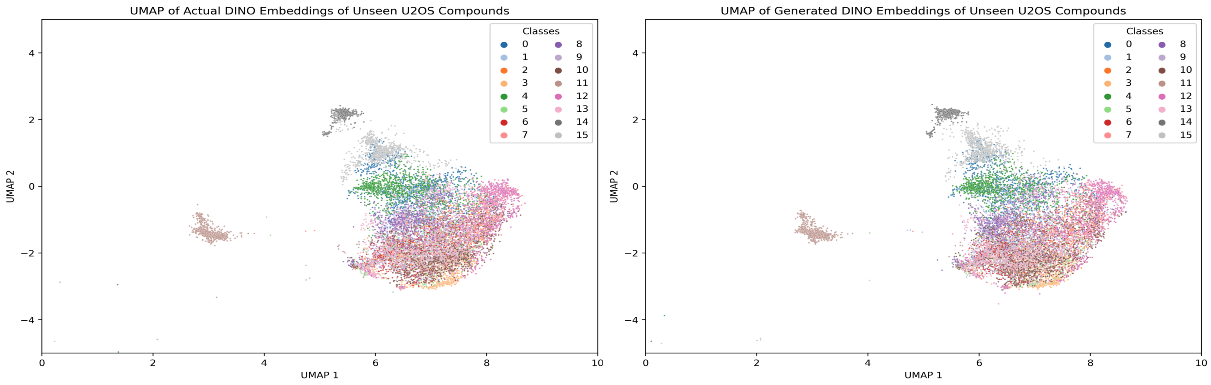


Figure A2: UMAP of real and generated embeddings for unseen genes on U2OS (JUMP)
